# Anniemap: Vector Search for Viral Short Read Alignment

**DOI:** 10.64898/2026.08.26.747390

**Authors:** Daniel J. van Zyl, Houriiyah Tegally, Cheryl Baxter, The INFORM Africa research study group, Tulio de Oliveira, Joicymara S. Xavier, Marcel Dunaiski

## Abstract

**Background:** The process of aligning sequencing reads to a reference genome is a foundational step in genomic analysis, underpinning tasks from variant detection to pathogen surveillance. In viral genomics, however, this problem becomes substantially more challenging: viral sequences are often present at low abundance within host-dominated samples and can differ markedly from available references due to rapid mutation and population heterogeneity. These characteristics reduce the effectiveness of conventional seed-and-extend aligners, which typically rely on long exact or near-exact matches to anchor alignments. Even modest sequence divergence or sequencing errors can disrupt such seeds, particularly for short reads, leading to missed alignments. The central challenge in this setting is maintaining robust alignment under high divergence without sacrificing efficiency.

**Results:** We introduce Anniemap, a vector search–based approach to viral short-read sequence alignment. Anniemap represents reads and reference sequences as binary vectors and performs approximate nearest-neighbour search using Facebook AI Similarity Search (FAISS) to efficiently identify candidate mappings. Anniemap was compared with the well-established alignment tools Bowtie2 and BWA-MEM2 across a diverse set of viral genomes and read lengths using both simulated and real sequencing data. Anniemap achieved higher sensitivity and throughput in almost all evaluated scenarios, with the most substantial improvements in sensitivity observed for highly divergent genomes, such as Hepatitis C virus (HCV) and Human Immunodeficiency Virus (HIV).

**Conclusions:** By measuring vector similarity rather than relying on long exact seed matches, Anniemap provides greater robustness to sequencing errors and genomic mutations. This property is particularly advantageous for viral genomes, where substantial sequence divergence is common. Further work is required to efficiently extend vector-based search for read alignment beyond viral genomes.

## Introduction

Viral genomics poses distinct challenges for read alignment, the task of assigning sequencing reads to their most likely genomic origin [1, 2]. In clinical and metagenomic settings, viral reads may be embedded within vast volumes of host-derived sequences [2], where even a small false positive rate can produce large numbers of spurious alignments. Further, sequence alignment becomes considerably more difficult when reads diverge from available references [2, 3], as is commonly the case for viral genomes, where high mutation rates and population heterogeneity can challenge even well-established alignment strategies [2, 4–6].

Next-generation sequencing technologies have dramatically reduced both the cost and time required to generate genomic data [7, 8]. Compared with long-read sequencing paradigms, short-read sequencing offers a practical combination of lower cost and higher per-base accuracy, making it well suited for population-scale studies and clinical variant discovery [9, 10]. These advantages are especially pronounced in viral genomics, where deep coverage and accurate base calling are essential for capturing rapidly evolving and highly diverse populations [3]. Within this domain, Illumina [11] sequencing has emerged as the dominant short-read platform [1, 7], routinely producing millions or billions of reads, typically 50–300 base pairs in length [2, 12]. The scale of these data imposes stringent computational demands which requires contemporary short read alingers to balance sensitivity, computational efficiency, and memory usage, while remaining robust to sequencing errors and genuine biological variation [6].

Many widely used short-read aligners use a seed-and-extend strategy [13–15] in which reads are decomposed into shorter subsequences (seeds) that are matched exactly or approximately against an indexed reference. Candidate mapping locations identified through these seed matches are then extended using dynamic programming or related alignment algorithms to produce full read alignments [14].

To achieve acceptable performance, many aligners rely on relatively long seeds to limit the number of candidate locations [6]. While larger seeds reduce computational cost, they are more likely to be disrupted by sequencing errors, point mutations, or indels, causing true alignments to be missed during the seeding phase [6, 16]. This problem is amplified in viral and other highly variable genomic contexts where mutations are frequent and broadly distributed. Consequently, aligners that depend on large exact or near-exact seeds may fail to detect valid read–reference relationships, motivating alternative approaches that tolerate greater sequence divergence during initial mapping [3, 16].

Vector search has recently been explored as an alternative to classical genomic sequence comparison, including taxonomic classification [17–19] and protein annotation [20]. Work on short-read alignment, however, remains limited. Embed-Search-Align (ESA) [21] maps transformer-based DNA embeddings against an online Pinecone vector store to align reads to the human genome, but reports a major throughput limitation: roughly 10,000 reads per minute versus roughly 1 million for Bowtie.

With this paper, we introduce Anniemap, a vector search–based framework for short-read viral sequence mapping. In Anniemap, sequences are represented as vectors and queried using approximate nearest-neighbour (ANN) search, an approach that has gained substantial prominence through its widespread adoption in large language models and other large-scale machine learning systems. Our framework makes use of FAISS (Facebook AI Similarity Search) [22], a high-performance ANN database architecture developed and publicly released by Meta. ANN search enables efficient retrieval of candidate mapping locations without exhaustive distance calculations, yielding run-times comparable to contemporary seed-and-extend aligners. By effectively considering a distribution of multiple short seeds rather than relying on longer individual seeds, Anniemap is more robust to sequencing errors and biological variation, particularly for shorter reads and highly diverse genomic populations. In contrast to ESA, we use simpler *k*-mer-based binary embeddings with an offline vector index to improve throughput, and we focus specifically on viral reads, which combine high mutation rates with comparatively short reference genomes. Anniemap achieved consistently higher sensitivity across all seven considered viruses, SARS-CoV-2, dengue (DENV), chikungunia (CHIKV), Ebola, Human Immunodeficiency Virus (HIV), Hepatitis C Virus (HCV) and influenza H1N1, compared to the two popular short read aligners Bowtie2 and BWA-MEM2, while also consistently achieving higher throughput than both.

## Methods

We reformulate the identification of candidate mappings as a vector search problem. By operating in a vector embedding space, this approach avoids reliance on long exact or near-exact matching seeds, with the goal of improving robustness to sequence divergence from the reference. The effectiveness of this formulation depends critically on the quality and discriminative power of the underlying vector representation. To enable efficient search at scale, we employ ANN search methods to identify candidate alignments. This allows for high-throughput querying with a controlled trade-off between sensitivity and computational cost relative to exhaustive search.

### Vector Representation

We represent sequences using a *k*-mer based vectorization scheme. While *k*-mer based representations provide a simple and efficient embedding for sequences, the vector search formulation introduced here is not tied to this choice. Alternative representations, including learned embeddings from deep neural networks or genomic language models, can be incorporated within the same paradigm.

### Binary *k*-mer

*k*-mer counting is a widely used technique in genomics, in which a sequence is represented by the frequency distribution of all substrings of length *k* drawn from an alphabet of size |Σ|, yielding a vector in |Σ|*^k^*-dimensional space. In Anniemap, rather than encoding *k*-mer counts, we construct binary vectors that indicate the presence or absence of each possible *k*-mer within a sequence.

This binary representation reduces both memory requirements and computational overhead. Although it discards frequency information, the resulting information loss is limited for short reads, where the probability of repeated *k*-mer within a single sequence is low and depends on both the choice of *k* and the sequence length [23]. Empirically, we found *k* = 5, which yields 1024-dimensional vectors, provides a favorable balance between discriminative power and computational efficiency.

### Canonicalization

To further improve efficiency, we provide an option to compute canonical *k*-mer, defined as the lexicographically smaller of a *k*-mer and its reverse complement. This reduces the effective dimensionality by a factor of two and ensures that forward and reverse complement sequences are represented identically, eliminating the need to handle strand orientation separately.

### Reference Sequence Indexing

Sequence mapping is performed relative to a collection of reference vector indexes derived from known reference sequences. This indexing strategy is conceptually analogous to seed-and-extend aligners, which also pre-index the reference genome to enable efficient candidate retrieval. However, in this case, the indexing is performed in a vector embedding space rather than over sequence seeds.

### Segmenting the Reference Sequence

For a fixed reference sequence *R* = (*r*_1_, *r*_2_, …, *r_N_*) of length *N* and a chosen read length *L_r_*, the reference database is constructed by segmenting *R* into overlapping subsequences of length *L_r_*:

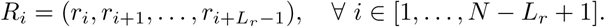

Each subsequence *R_i_* is independently transformed into a binary *k*-mer vector and inserted into the vector database. This procedure yields approximately *N* − *L_r_* + 1 reference vectors, providing dense coverage of the reference sequence. By default, we generate subsequences with a stride of one base pair to maximize sensitivity. A larger stride reduces the database size and improves throughput, but may decrease mapping resolution and sensitivity.

To accommodate the variable read lengths produced by Illumina sequencing, separate reference indices may be constructed for a range of read lengths. For a query read of length *L_q_*, we select the index corresponding to the largest reference read length *L_r_* ≤ *L_q_*.

For canonical vector representations, index construction stores both the binary *k*-mer presence vectors as well as a corresponding strand vector

The strand vector encodes orientation per canonical *k*-mer. For a read represented over *d* canonical *k*-mers, the strand vector *s* ∈ {0, 1}*^d^* is defined as

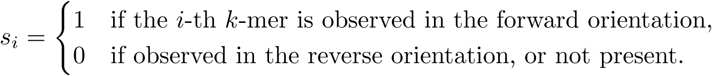

The strand vector is retained for downstream reconstruction of query strand information.

For the strand vectors, we only consider the first occurrence of any repeating *k*-mers.

### Inverted File Index

For efficient similarity search over large collections of fixed-dimensional binary vectors, we make use of FAISS [22]. Specifically, we leverage the Binary Inverted File (IVF) index.

Let 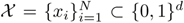 denote a collection of *N* binary vectors of dimension *d*. In our case, these are our reference vectors. The IVF framework partitions *X* into *n*_list_ clusters, each represented by a centroid 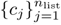.

The similarity between the vectors is measured using Hamming distance. For *x, y* ∈ {0, 1}*^d^*, this is defined as

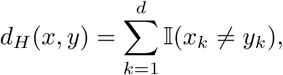

where I(·) is the indicator function. Each vector is then assigned to its nearest centroid under this metric,

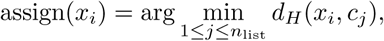

and stored in the corresponding inverted list. The resulting index consists of the centroids together with their associated vector lists.

### Vector Search

Given a query vector *q* ∈ {0, 1}*^d^*, representing a read encoded as a binary vector, the same Hamming distance is used to identify the *n*_probe_ nearest centroids. The search is then restricted to the inverted lists associated with these centroids, significantly reducing the number of candidate vectors from *N* to a much smaller subset.

The parameter *n*_list_ thus controls the granularity of the partitioning, while *n*_probe_ determines how many clusters are explored at query time. Increasing *n*_probe_ typically improves recall by enlarging the search space, at the cost of higher query latency.

For canonical vector representations, strand assignment is performed after vector retrieval. For each retrieved candidate, the corresponding reference strand vector is obtained from a precomputed strand table.

Let *q*^(*s*)^, *r*^(*s*)^ ∈ {0, 1}*^d^* denote the strand bit-vectors of the query and reference, and let *q*^(*p*)^, *r*^(*p*)^ ∈ {0, 1}*^d^* denote their presence vectors. Strand comparison is restricted to overlapping canonical *k*-mers, defined by the mask

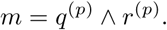

We sum strand agreement over the overlapping positions and assign the mapping strand by majority vote.

### Sequence Alignment

After identifying candidate alignment locations, we perform dynamic programming-based sequence alignment similar to seed-and-extend aligners. Specifically, we employ the Wavefront Alignment (WFA2) algorithm [24, 25], an exact algorithm for gap-affine alignment that leverages homologous regions between sequences to accelerate computation.

Given two sequences of lengths *n* and *m*, respectively, traditional dynamic programming approaches compute optimal alignments under a gap-affine scoring model with quadratic time complexity. In contrast, the WFA algorithm computes exact alignments with time complexity

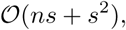

where *s* is the alignment score, and uses *O*(*s*^2^) memory.

### Scoring Scheme

Anniemap uses a gap-affine scoring model (Smith–Waterman–Gotoh). This scoring includes a mismatch penalty *X*, and gaps of length *ℓ* are penalized according to

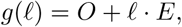

where *O* is the gap opening penalty and *E* is the gap extension penalty. This formulation favors longer contiguous gaps over multiple short gaps, which is appropriate in many genomic alignment settings. A similar schoring scheme is used by both Bowtie2 and BWA-MEM2.

### Heuristics

By default, the WFA algorithm guarantees recovery of the optimal alignment under the specified scoring model. To improve computational throughput, we additionally employ a static banded heuristic and the Z-drop criterion.

The banded heuristic restricts the search space to a fixed diagonal interval, preventing the wavefront from expanding beyond predefined limits. Specifically, alignment is constrained to diagonals *k* ∈ [*k*_min_, *k*_max_], thereby reducing computational overhead while preserving alignments within the allowed band.

The Z-drop heuristic, as described in [26], terminates extension when the alignment score drops sharply relative to the best score observed so far, while accounting for diagonal displacement between alignment paths.

### Viral Read Simulation

In the absence of ground-truth mappings for real sequencing data, we evaluated Anniemap using simulated paired-end short-read datasets generated from high-quality viral consensus genomes obtained from NCBI. Simulation provides precise control over read characteristics, including length and sequencing error profiles, while also enabling exact ground-truth alignment for quantitative evaluation.

We considered a set of clinically relevant viruses, including SARS-CoV-2, dengue virus (all four serotypes), chikungunya virus (Asian and ECSA lineages), HIV, Influenza A H1N1 (all eight segments), Hepatitis C virus (genotypes 1, 2, 3, 4, and 6), and Ebola virus. Furthermore, we only included complete genomes with greater than 97.5% coverage and no ambiguous nucleotides.

For each virus, we sampled multiple distinct reference genomes to capture genomic diversity, namely, 100 genomes each for SARS-CoV-2, Ebola, and HIV; 50 genomes for each chikungunya lineage; and 25 genomes for each of the remaining virus groups. From each genome, 25,000 paired-end reads were simulated using Mason2 [27]. We conducted experiments across read lengths of 50, 75, 100, 150 and 300 base pairs.

Our read simulation used the default Mason2 error model, with the mismatch scaling factor set to 2.5×, consistent with the configuration used to evaluate Minimap2 [26]. To further assess robustness under extreme divergence, we conducted an additional experiment using an increased mismatch scaling factor of 25×.

To evaluate specificity in a realistic background setting, we also simulated 25 million paired-end reads from the human reference genome (GRCh38) to represent host-derived sequences and to quantify false positive alignments under high-background conditions.

### Parameter Tuning

For Anniemap, we expose several parameters that control both indexing and alignment. These include the IVF parameters *n*_list_ and *n*_probe_, the *k*-mer length *k*, and alignment-related parameters such as mismatch penalty, gap opening and extension penalties, Z-drop threshold, and minimum alignment score.

Parameter tuning was performed via grid search across the different stages of the alignment pipeline, using an independent validation set comprising 2.5 million simulated paired-end reads each from SARS-CoV-2, dengue serotype 1, and HIV. These viruses were selected to span a range of sequence divergence, with SARS-CoV-2 representing a relatively conserved genome and HIV representing a highly diverse one. The objective was not exhaustive optimization, but rather the selection of a reasonable parameter configuration across these representative cases that could subsequently be evaluated for generalizability on the independent test set.

The alignment score threshold was calibrated using 25 million simulated paired-end human reads aligned against SARS-CoV-2, dengue serotype 1, and HIV references, with the aim of matching the false positive rates of Bowtie2 and BWA-MEM2. Under this setting, no false positives were observed for dengue or HIV. For SARS-CoV-2, a small number of false positives (0.3%) were observed at 50 bp, attributable to spurious long homologous stretches of low-complexity sequence within the reference.

## Results & Discussion

All results are reported on an independent test set. For chikungunya, dengue, and HCV, recall is aggregated across all genotypes. For Influenza A H1N1, recall is aggregated across all eight genome segments; the reference index includes all segments jointly rather than treating them independently. Bowtie2 and BWA-MEM2 were run using their default parameters.

For 300 base pair reads, Anniemap performs the vector search using only the central 150 base pairs, as increasing the vector density beyond this length results in excessive binary collisions within the 512-dimensional vector embeddings. The subsequent alignment is nevertheless performed across the full read length.

### Alignment Sensitivity

Additional file 1 contains the recall scores of Bowtie2, BWA-MEM and Anniemap over the different considered viruses across all read lengths. Figure 1 offers a visual representation of the same data.

**Fig. 1.**
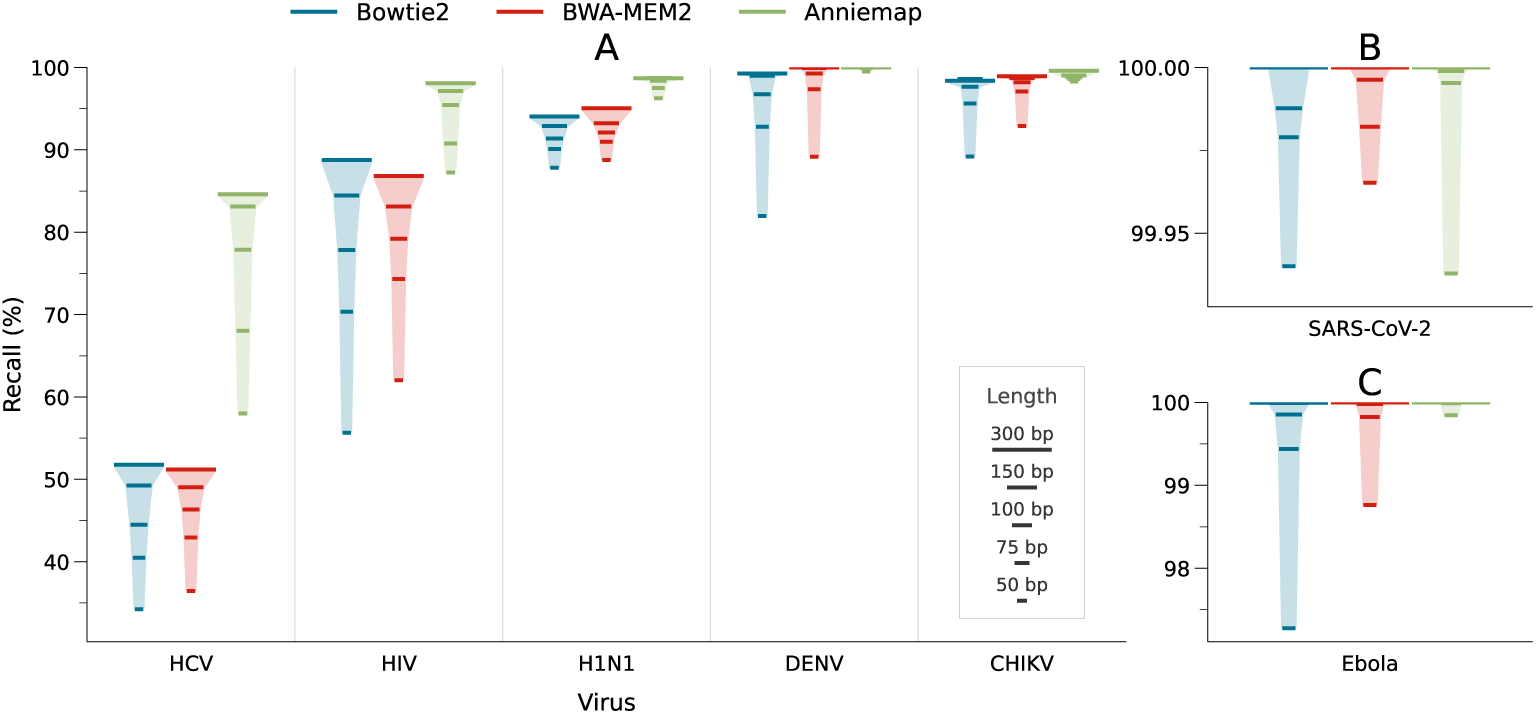
Comparison of recall for Bowtie2 (blue), BWA-MEM2 (red), and Anniemap (green) across multiple viruses and read lengths under a standard error model (2.5× mismatch scaling). Horizontal bars correspond to read lengths of 50, 75, 100, 150 and 300 base pairs (in increasing order, with proportional widths). Panels B and C show recall for SARS-CoV-2 and Ebola, respectively, with axes scaled independently to better capture differences in performance.

For the purposes of these experiments, a read is considered correctly mapped if its longest alignment overlaps the true interval, with an overlap of at least 50% of the true interval length.

Across the experiments, Anniemap consistently achieves the highest recall, with the most pronounced gains observed for viruses that exhibit greater sequence divergence. Figure 2 shows the per site nucleotide divergence distribution for each virus. For relatively conserved genomes such as SARS-CoV-2 and Ebola, all methods perform near perfectly, with recall approaching 100% even at shorter read lengths. The SARS-CoV-2 results in particular reflect a limited challenge posed by low-divergence sequences where differences between methods are negligible. The 50 base pair read length experiment for SARS-CoV-2 was the only condition under which Anniemap exhibited lower sensitivity than the other tools.

**Fig. 2.**
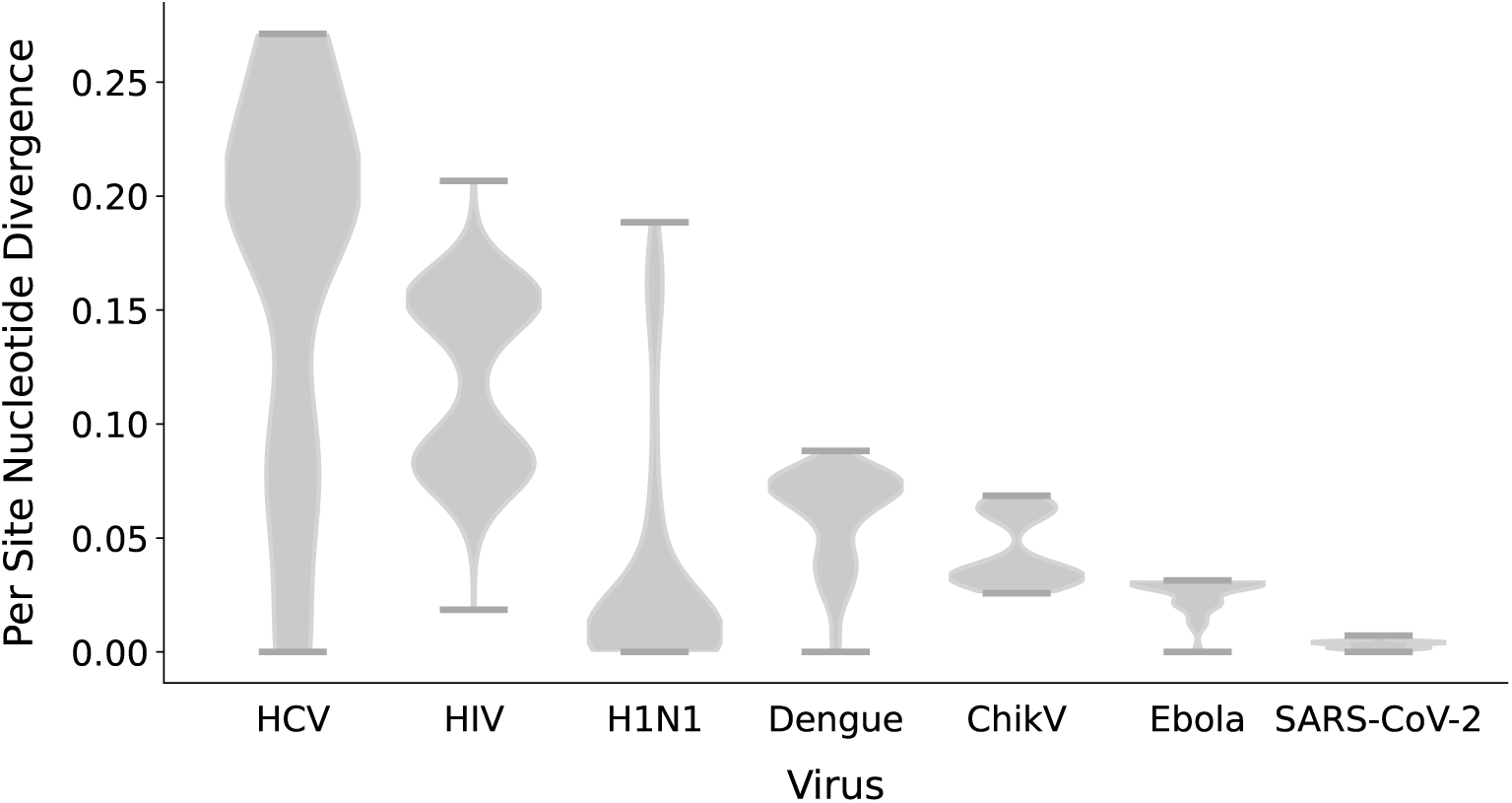
Per-site nucleotide divergence across viral genomes, computed as the total number of substitutions, insertions, and deletions relative to a reference sequence, normalized by the length of the aligned reference.

In contrast, more substantial differences emerge for viruses with higher genetic diversity. For dengue and chikungunya, Anniemap maintains near-perfect recall across all read lengths, while Bowtie2 and BWA-MEM2 show reduced sensitivity, particularly at shorter read lengths. Although this gap narrows as read length increases, Anniemap retains a consistent, albeit smaller, advantage across all read lengths.

The performance differences between compared tools is most pronounced for highly variable viruses such as HIV and HCV. For HIV, Anniemap achieves notably higher recall regardless of read length. HCV represents the most challenging case in this benchmark, where all methods experience significantly reduced recall.

Influenza A H1N1 constitutes an intermediate case, where Anniemap again outperforms the competing methods across all read lengths, although the margin is smaller than for HIV and HCV.

These trends are consistent with the reliance of seed-and-extend aligners such as Bowtie2 and BWA-MEM2 on exact or near-exact seed matches. Shorter reads reduce the likelihood of observing sufficiently long exact matches, particularly in the presence of sequence divergence. In contrast, by avoiding dependence on long exact seeds, Anniemap maintains higher sensitivity across a range of read lengths, with the largest gains observed in the most divergent viral populations.

### Increasing Sequence Error Rate

As an additional experiment, we consider an extreme divergence scenario in which the sequencing error rate is artificially increased using Mason2 to stress-test robustness to sequence divergence. This setting is not intended to reflect realistic conditions for the viruses considered, but rather to evaluate performance under highly degraded sequence similarity.

Additional file 1 contains the recall scores of Bowtie2, BWA-MEM and Anniemap over the different considered viruses across all read lengths under these conditions.

Under the extreme error setting (25× mismatch scale), recall decreases for all methods, with the extent of degradation varying by virus and read length (Figure 3). Despite this, Anniemap consistently maintains substantially higher recall than both Bowtie2 and BWA-MEM2 under all conditions.

**Fig. 3.**
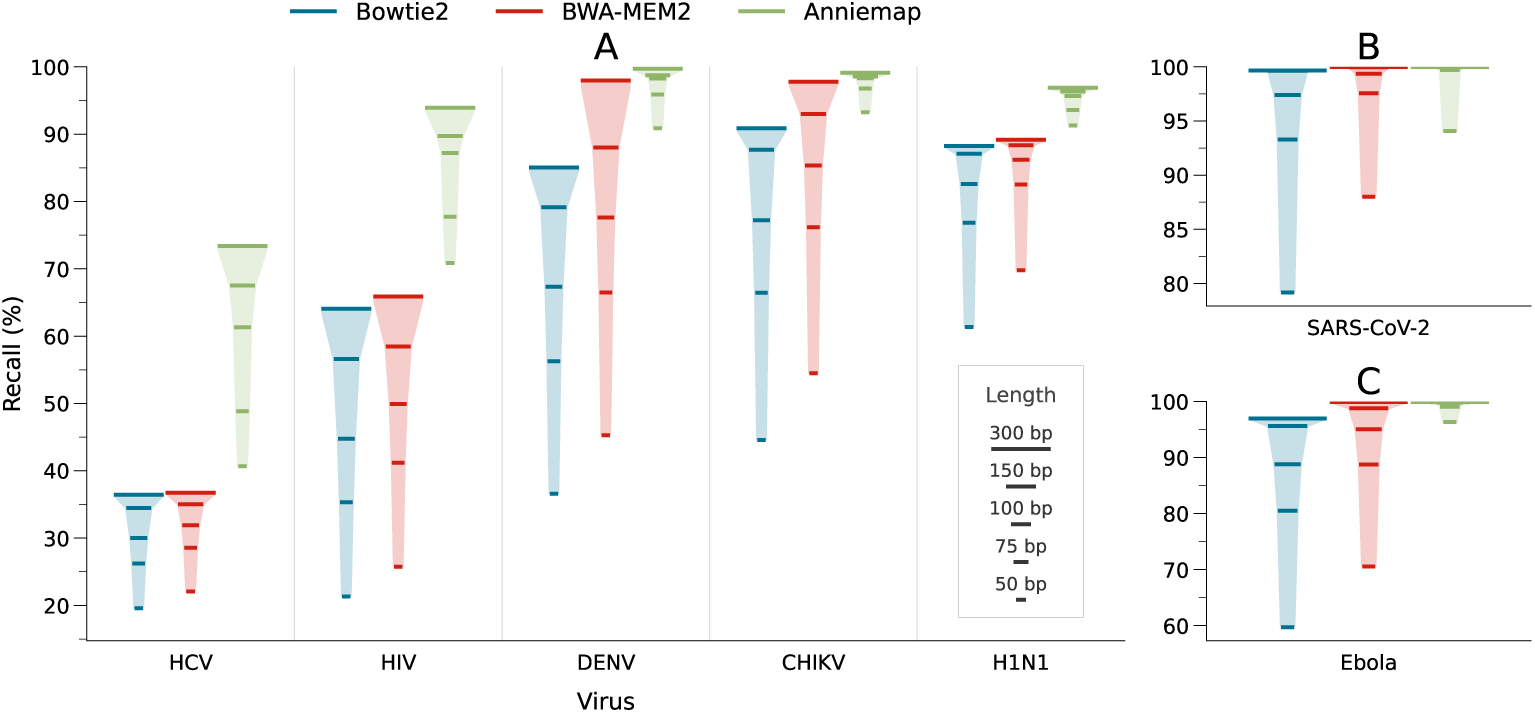
Comparison of recall for Bowtie2 (blue), BWA-MEM2 (red), and Anniemap (green) across multiple viruses and read lengths under an extreme error model (25× mismatch scaling). Horizontal bars correspond to read lengths of 50, 75, 100, and 150 base pairs (in increasing order, with proportional widths). Panels B and C show recall for SARS-CoV-2 and Ebola, respectively, with axes scaled independently to better capture differences in performance.

The largest relative performance gaps are observed for viruses with moderate underlying sequence divergence. Although Anniemap exhibits a reduction in sensitivity under increased error rates, the decline is markedly smaller than that observed for Bowtie2 and BWA-MEM2.

For highly diverse viruses such as HIV and HCV, the additional error burden results in comparatively smaller relative decreases in recall, as baseline performance is already constrained by divergence.

For more conserved viruses, including SARS-CoV-2 and Ebola, performance differences become more apparent under elevated error rates, with Anniemap achieving the highest recall. Notably, even under this extreme setting, Anniemap attains 99.091% recall for Ebola and 99.707% for SARS-CoV-2 at 75 bp read lengths.

Among all viruses considered, the Influenza A H1N1 sensitivity results were least affected by the increased error rate. This is likely attributable to the relatively short length of its genome segments.

### Real Data Experiments

To complement the simulated data experiments, we conducted an additional evaluation using real Illumina sequencing data obtained from the Sequence Read Archive (SRA). The analysis included reads from three BioProjects submitted by the KwaZulu-Natal Research Innovation and Sequencing Platform (KRISP): PRJNA1231728 for dengue virus serotype 1, PRJNA772074 for SARS-CoV-2, and PRJNA1197182 for HIV. These datasets correspond to the same three viruses used in our simulated validation experiments and collectively provide a representative range of viral sequence diversity.

For each virus, a collection of samples was obtained and filtered to exclude samples in which fewer than 50% of reads were identified as viral by all three evaluated tools. The datasets contained a range of read lengths; however, reads shorter than 50 base pairs were excluded from further analysis. From the remaining data, we selected 10 dengue samples (18,410,502 viral reads), 50 SARS-CoV-2 samples (18,439,231 viral reads), and 200 HIV samples (22,429,158 viral reads).

In contrast to the simulated datasets, no ground-truth alignments are available for the real sequencing data, which prevents a direct assessment of mapping correctness. Instead, we compared the proportion of reads successfully mapped by each tool and examined differences in mapping behaviour across datasets.

Figure 4 shows the proportion of reads mapped by each tool relative to the total number of reads mapped by at least one of the three methods. For the HIV and dengue datasets, Anniemap mapped a larger proportion of reads than both Bowtie2 and BWA-MEM2, consistent with the trends observed in the simulated experiments. In contrast, the SARS-CoV-2 results indicate that all three tools mapped a highly similar proportion of reads. The relative differences between tools were not identical to those observed in the simulated experiments; however, the real datasets were derived exclusively from samples representing the genetic diversity present within Southern Africa.

**Fig. 4.**
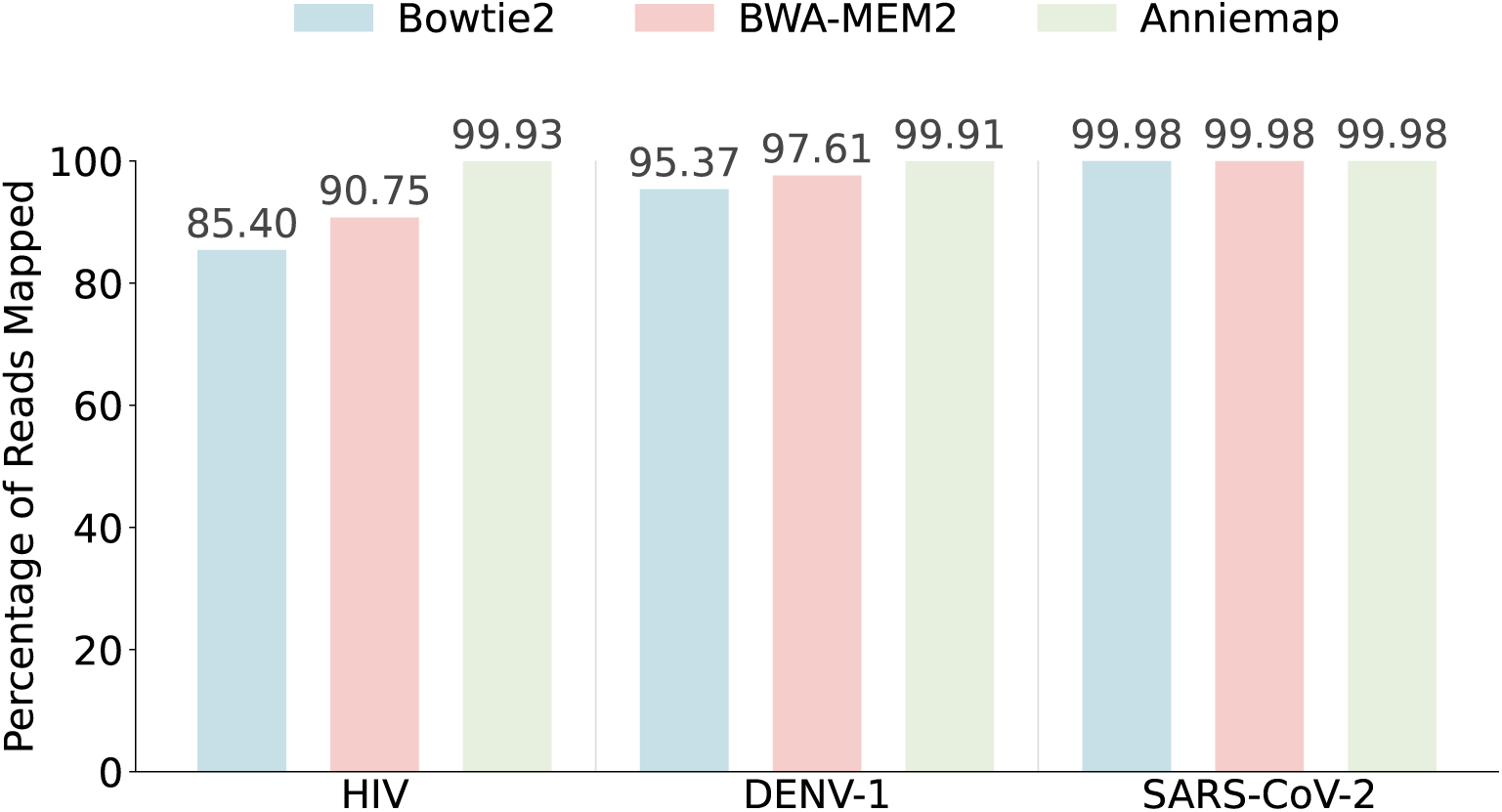
Comparison of the proportion of reads mapped by Bowtie2 (blue), BWA-MEM2 (red), and Anniemap (green) among all reads mapped by at least one of the evaluated tools. Reads were considered successfully mapped only if the aligned segment covered at least 50% of the original read length, thereby excluding excessively short alignments likely to represent spurious matches.

Although Anniemap consistently mapped more reads than Bowtie2 and BWA-MEM2, we cannot verify the mapping correctness. Nevertheless, the unmapped reads from Bowtie2 and BWA-MEM2 can be analysed to identify the potential causes of mapping failure. Of particular concern would be if Anniemap mapped additional reads solely because it accepted alignments with lower alignment scores, implying a more permissive alignment threshold. However, Table 1 indicates that this was not the dominant factor.

**Table 1.** Breakdown of reads exclusively mapped by Anniemap across the HIV, dengue, and SARS-CoV-2 datasets. For each virus, the values indicate the percentage of Anniemap-exclusive reads for which Bowtie2 or BWA-MEM2 failed to report a mapping due to the specified cause category, including failure to identify a candidate mapping location, identification of an alternative candidate location, insufficient alignment score, or alignment length below the acceptance threshold.

| **Virus** | **Category** | **Bowtie2** | **BWA-MEM2** |
| --- | --- | --- | --- |
| **HIV** (n=1,599,690) | No Candidate Mapping | 96.67 | 91.66 |
|  | Different Alignment Locations | 2.69 | 4.59 |
|  | Alignment Score Too Low | 0.13 | 0.06 |
|  | Alignment Too Short | 0.51 | 3.69 |
| **DENV** (n=425,195) | No Candidate Mapping | 86.50 | 85.26 |
|  | Different Alignment Locations | 10.56 | 11.47 |
|  | Alignment Score Too Low | 0.04 | 0.00 |
|  | Alignment Too Short | 2.90 | 3.27 |
| **SARS-CoV-2** (n=2,700) | No Candidate Mapping | 2.30 | 1.48 |
|  | Different Alignment Locations | 50.48 | 51.52 |
|  | Alignment Score Too Low | 0.07 | 0.04 |
|  | Alignment Too Short | 47.15 | 46.96 |

For reads mapped by Anniemap but not by both Bowtie2 and BWA-MEM2, the predominant reason for a missed mapping in the dengue and HIV datasets was the inability of the latter tools to identify a candidate mapping location. This finding is consistent with both the simulated experiments and the theoretical motivation underlying Anniemap’s design, namely the avoidance of strict long exact-match seed requirements through the use of vector similarity methods. Candidate mapping therefore represents the principal distinction between Anniemap and the other evaluated tools, whereas all three methods employ broadly similar Smith–Waterman-based alignment scoring approaches.

The generally high level of agreement between the three tools further suggests that, when Bowtie2 and BWA-MEM2 were able to identify appropriate candidate mapping locations, their final alignments were often consistent with those produced by Anniemap. The remaining cases in which Bowtie2 and BWA-MEM2 failed to map reads successfully mapped by Anniemap were primarily attributable to candidate mappings being identified at differing reference locations, resulting in poor downstream alignment scores, or to the resulting alignments being considered too short.

These latter failure modes were observed more frequently in the SARS-CoV-2 datasets, for which missed candidate mappings would be less expected given the comparatively high sequence conservation of SARS-CoV-2. Consequently, the proportion of reads mapped exclusively by Anniemap was substantially lower for SARS-CoV-2 (0.014%) than for dengue (1.860%) or HIV (7.091%). Across all analysed datasets, there were very few cases in which Bowtie2 or BWA-MEM2 failed to report an alignment due to the alignment score falling below the acceptance threshold.

These findings further indicate that Anniemap is able to identify more suitable candidate mapping locations than Bowtie2 and BWA-MEM2 for divergent viral reads.

### Throughput and Memory Usage

We measured the total runtime of each tool on the hold-out test set for each virus. The results show a consistent trend across all datasets (Figure 5). At shorter read lengths, all three tools exhibit comparable runtimes. However, as read length increases, Anniemap maintains relatively stable runtime, while Bowtie2 shows the largest increase. This behaviour is expected, as Anniemap’s vector representations have fixed dimensionality and do not scale with read length. In contrast, seed-and-extend approaches incur increased computational cost as read length increases.

**Fig. 5.**
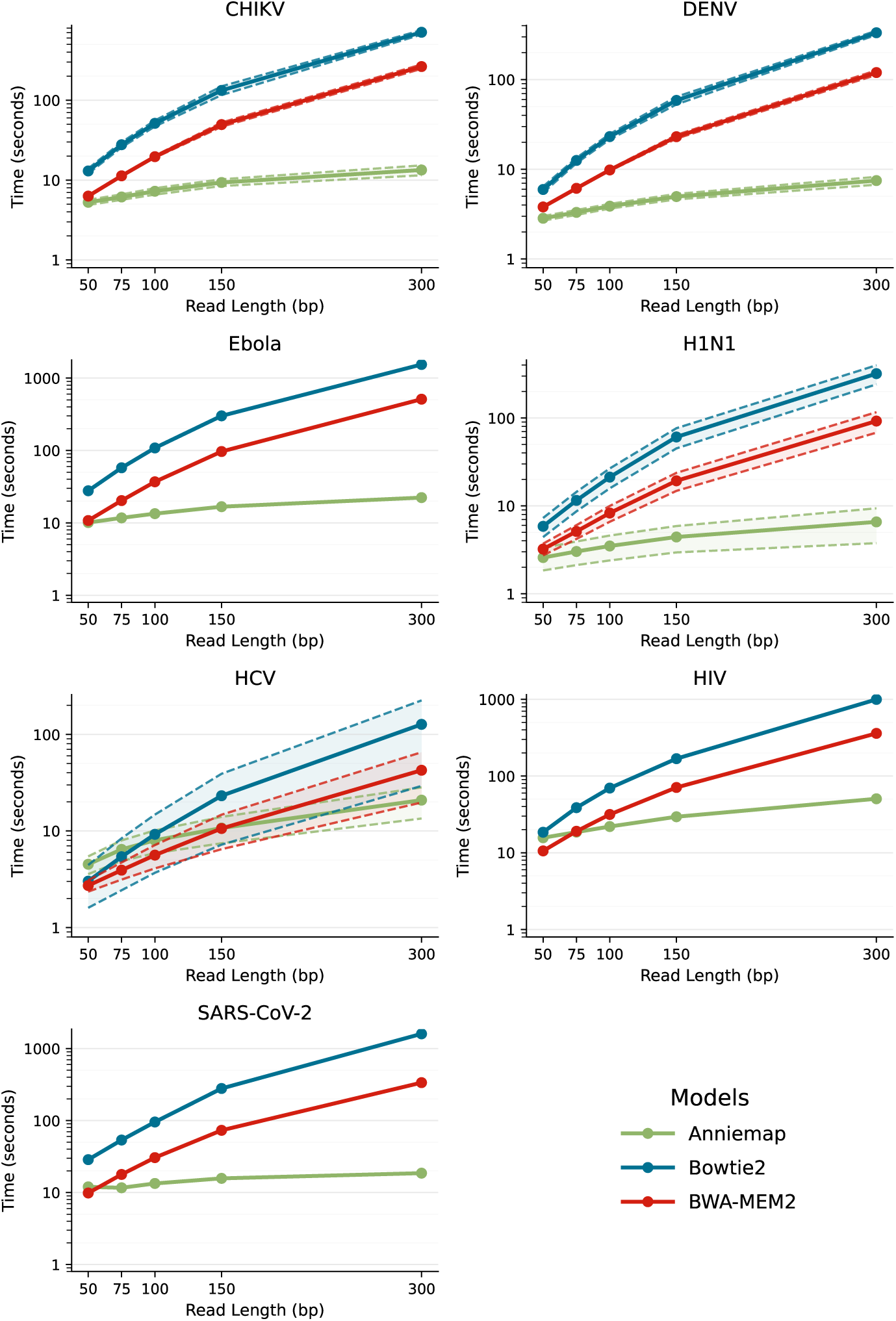
Comparison of runtime for Bowtie2 (blue), BWA-MEM2 (red), and Anniemap (green) across different viruses and read lengths at a standard error rate. The shaded regions, bounded by dashed lines, indicate the standard deviation over five independent runs for each model. All tools were executed using four threads.

For HCV and HIV, Anniemap exhibits a reduced runtime advantage compared to Bowtie2 and BWA-MEM2, with BWA-MEM2 performing best on HCV. This is likely attributable to the lower recall of Bowtie2 and BWA-MEM2 on these viruses under higher sequence divergence, which reduces the number of candidate seeds identified. Consequently, fewer downstream alignment operations are performed, which results in comparatively lower observed runtimes for these tools.

We further evaluated the tools in terms of peak memory usage, with the results presented in Table 2. Bowtie2 consistently exhibited the lowest peak memory usage across all evaluated conditions, while Anniemap used marginally more memory. BWA-MEM2 exhibited substantially higher memory usage than both other tools. As expected, peak memory usage increased across all three tools with increasing read length.

**Table 2.** Peak memory usage (Megabytes) of Bowtie2, BWA-MEM2 and Anniemap for different viruses across various read lengths at a standard error rate (2.5x mismatch scale).

| Virus | Bowtie2 | BWA-MEM2 | Anniemap |
| --- | --- | --- | --- |
| **50bp** |  |  |  |
| CHIKV | 51.5 | 807.5 | 80.1 |
| DENV | 51.3 | 644.9 | 78.7 |
| Ebola | 51.4 | 955.2 | 83.8 |
| H1N1 | 51.4 | 664.7 | 83.2 |
| HCV | 53.6 | 564.1 | 78.5 |
| HIV | 51.8 | 894.5 | 82.6 |
| SARS-CoV-2 | 51.8 | 1217.7 | 85.8 |
| **75bp** |  |  |  |
| CHIKV | 58.5 | 709.6 | 87.9 |
| DENV | 58.3 | 651.0 | 88.7 |
| Ebola | 58.3 | 844.7 | 94.5 |
| H1N1 | 58.3 | 615.6 | 93.0 |
| HCV | 60.4 | 525.4 | 89.1 |
| HIV | 58.6 | 787.0 | 93.5 |
| SARS-CoV-2 | 58.9 | 858.9 | 94.0 |
| **100bp** |  |  |  |
| CHIKV | 64.5 | 666.2 | 97.6 |
| DENV | 64.0 | 589.3 | 99.3 |
| Ebola | 64.1 | 835.8 | 101.2 |
| H1N1 | 64.0 | 607.5 | 104.3 |
| HCV | 66.1 | 514.6 | 101.4 |
| HIV | 64.6 | 735.5 | 100.3 |
| SARS-CoV-2 | 64.7 | 829.0 | 106.5 |
| **150bp** |  |  |  |
| CHIKV | 91.6 | 608.9 | 124.1 |
| DENV | 90.4 | 530.6 | 125.5 |
| Ebola | 90.2 | 744.3 | 125.8 |
| H1N1 | 90.4 | 528.9 | 131.6 |
| HCV | 92.1 | 476.7 | 130.7 |
| HIV | 92.0 | 710.0 | 128.9 |
| SARS-CoV-2 | 91.7 | 750.8 | 135.8 |
| **300bp** |  |  |  |
| CHIKV | 161.7 | 521.4 | 210.3 |
| DENV | 153.3 | 526.9 | 225.2 |
| Ebola | 143.7 | 538.8 | 224.7 |
| H1N1 | 157.1 | 521.9 | 236.2 |
| HCV | 167.4 | 463.5 | 223.1 |
| HIV | 166.7 | 609.7 | 215.7 |
| SARS-CoV-2 | 145.9 | 613.5 | 218.5 |

## Conclusion

Reformulating candidate mapping as a vector similarity search problem removes the reliance on long exact or near-exact matching seeds that underpin conventional aligners. This shift enables improved robustness in settings where sequence similarity to available references is degraded, whether due to biological divergence or sequencing error. The benefits of this approach are most evident for genetically diverse viral populations, where traditional seeding strategies are more prone to failure. More broadly, this increased tolerance to sequence deviation may be advantageous in scenarios where coverage is incomplete or sequence quality is compromised, including challenging environments such as wastewater sequencing, where viral abundance is low and sequences may be substantially degraded.

We evaluated our vector search across a range of viral genomes and sequencing conditions. Anniemap consistently demonstrates improved sensitivity relative to established aligners such as Bowtie2 and BWA-MEM2, particularly in regimes of increased divergence and shorter read lengths. By employing an inverted file (IVF) index for vector search, Anniemap achieves this improved robustness while consistently maintaining higher throughput than the compared tools.

An important limitation of Anniemap is its current applicability beyond RNA viral genomes. The sensitivity advantages observed over traditional aligners are less pronounced for longer genomes with lower mutation rates. Furthermore, as reference genome length increases, Anniemap, without further optimisation, experiences a substantially greater reduction in throughput compared with Bowtie2 and BWA-MEM2.

While these results are promising, Anniemap should be viewed as an proof of concept within a broader promising paradigm. The vector search formulation decouples sequence representation from the search procedure, creating opportunities for future improvement. In particular, more expressive embedding strategies—potentially derived from deep learning models or genomic language models may further enhance sensitivity and application. The computational structure of vector search is also well-suited to hardware acceleration, with the potential of further throughput gains.

## Declarations

### Competing interests

No competing interest is declared.

### Availability of data and materials

Source code is available at https://github.com/INFORM-Africa/AI-viral-lineage-classification.

The NCBI Sequences used in this research are openly available and can be downloaded from https://www.ncbi.nlm.nih.gov/labs/virus/vssi/. The SRA Reads used in this research are openly available and can be downloaded from https://www.ncbi.nlm.nih.gov/sra.

Scripts to download the exact sequence datasets used from these sequence archives are provided at at https://github.com/INFORM-Africa/AI-viral-lineage-classification.

### Funding

Research activities were supported in part by grants from the National Institute of Health, INFORM Africa project through IHVN (U54 TW012041), as well as the UK Medical Research Foundation (MRF-RG-ICCH-2022-100069), the Wellcome Trust through the Global.health project (228186/Z/23/Z), and the Novo Nordisk Foundation (NNF24OC0094346).

### Authors’ contribution

DJvZ, JSX, MD, HT, CB, and TdO devised the project. DJvZ developed the models, performed all the experiments, and wrote the paper. MD, JSX, TdO, and CB manage the project and funding. The INFORM Africa research study group supported the project implementation, and all authors read and reviewed it.

### INFORM Africa Research Study Group for D-SI Africa Consortium

Akros: Christina Riley, Anna Winters. Centre for the AIDS Programme of Research in South Africa (CAPRISA): Vivek Naranbhai, Felix Made, Salim Abdool Karim. Consortium for Advanced Research Training in Africa (CARTA): Kennedy Otwombe. Institute of Human Virology Nigeria: Alash’le Abimiku, Sophia Osawe, James Onyemata, Patrick Dakum, Fati Murtala-Ibrahim, Nifarta Andrew, Aminu Musa, Tolulope Adenekan, Kenneth Ewerem, Victoria Etuk. Stellenbosch University/Centre for Epidemic Response and Innovation (CERI):Tulio de Oliveira, Cheryl Baxter, Eduan Wilkinson, Houriiyah Tegally, Jenicca Poongavanan, Michelle Parker, Danilo Silva, Joicymara S. Xavier. University of Maryland Baltimore: Kristen A. Stafford, Manhattan Charurat, Natalia Blanco, Timothy O’Connor, Meagan Fitzpatrick, Mohammad M. Sajadi. University of Port Harcourt. Olanrewaju Lawal. Villanova University: Chenfeng Xiong, Weiyu Luo, Xin Wu.

### Competing interests

No competing interest is declared.

### Ethics approval and consent to participate

Not applicable

### Consent for publication

Not applicable

## Notes

### Competing Interest Statement

The authors have declared no competing interest.

## References

[1] Hu T, Chitnis N, Monos D, Dinh A. Next-generation sequencing technologies: An overview. Human Immunology. 2021;82(11):801–811. Next Generation Sequencing and its Application to Medical Laboratory Immunology. 10.1016/j.humimm.2021.02.012.

[2] Rose R, Constantinides B, Tapinos A, Robertson DL, Prosperi M. Challenges in the analysis of viral metagenomes. Virus Evolution. 2016 08;2(2):vew022. 10.1093/ve/vew022. https://academic.oup.com/ve/article-pdf/2/2/vew022/24139230/vew022.pdf.

[3] McElroy K, Thomas T, Luciani F. Deep sequencing of evolving pathogen populations: applications, errors, and bioinformatic solutions. Microbial Informatics and Experimentation. 2014;4(1):1. 10.1186/2042-5783-4-1.

[4] Zielezinski A, Vinga S, Almeida J, Karlowski W. Alignment-free sequence comparison: benefits, applications, and tools. Genome Biology. 2017;18(1):186. 10.1186/s13059-017-1319-7.

[5] Wang Q, Jia P, Zhao Z. VERSE: a novel approach to detect virus integration in host genomes through reference genome customization. Genome Medicine. 2015;7(1):2. 10.1186/s13073-015-0126-6.

[6] Alser M, Rotman J, Deshpande D, Taraszka K, Shi H, Baykal PI, et al. Technology Dictates Algorithms: Recent Developments in Read Alignment. Genome Biology. 2021;22(1):249. 10.1186/s13059-021-02443-7.

[7] Goodwin S, McPherson JD, McCombie WR. Coming of age: ten years of next-generation sequencing technologies. Nature Reviews Genetics. 2016;17(6):333–351. 10.1038/nrg.2016.49.

[8] Schadt EE, Turner S, Kasarskis A. A window into third-generation sequencing. Human Molecular Genetics. 2010 09;19(R2):R227–R240. 10.1093/hmg/ddq416. https://academic.oup.com/hmg/articlepdf/19/R2/R227/1798881/ddq416.pdf.

[9] Polonis K, Blommel JH, Hughes AEO, Spencer D, Thompson JA, Schroeder MC. Innovations in Short-Read Sequencing Technologies and Their Applications to Clinical Genomics. Clinical Chemistry. 2025 01;71(1):97–108. 10.1093/clinchem/hvae173. https://academic.oup.com/clinchem/articlepdf/71/1/97/61283954/hvae173.pdf.

[10] Watson SJ, Welkers MRA, Depledge DP, Coulter E, Breuer JM, de Jong MD, et al. Viral population analysis and minority-variant detection using short read next-generation sequencing. Philosophical Transactions of the Royal Society B: Biological Sciences. 2013 03;368(1614):20120205. 10.1098/rstb.2012.0205. https://royalsocietypublishing.org/rstb/articlepdf/doi/10.1098/rstb.2012.0205/253257/rstb.2012.0205.pdf.

[11] Bentley DR, Balasubramanian S, Swerdlow HP, Smith GP, Milton J, Brown CG, et al. Accurate whole human genome sequencing using reversible terminator chemistry. Nature. 2008;456(7218):53–59. 10.1038/nature07517.

[12] Buermans HPJ, den Dunnen JT. Next generation sequencing technology: Advances and applications. Biochimica et Biophysica Acta (BBA) - Molecular Basis of Disease. 2014;1842(10):1932–1941. From genome to function. 10.1016/j.bbadis.2014.06.015.

[13] Roberts M, Hayes W, Hunt BR, Mount SM, Yorke JA. Reducing storage requirements for biological sequence comparison. Bioinformatics. 2004 07;20(18):3363–3369. 10.1093/bioinformatics/bth408.

[14] Yan Y, Chaturvedi N, Appuswamy R. Accel-Align: a fast sequence mapper and aligner based on the seed–embed–extend method. BMC Bioinformatics. 2021;22:257. 10.1186/s12859-021-04162-z.

[15] Ye H, Meehan J, Tong W, Hong H. Alignment of Short Reads: A Crucial Step for Application of Next-Generation Sequencing Data in Precision Medicine. Pharmaceutics. 2015;7(4):523–541. 10.3390/pharmaceutics7040523.

[16] Hoffmann S, Otto C, Kurtz S, Sharma CM, Khaitovich P, Vogel J, et al. Fast mapping of short sequences with mismatches, insertions and deletions using index structures. PLoS Computational Biology. 2009;5(9):e1000502. Epub 2009 Sep 11. 10.1371/journal.pcbi.1000502.

[17] Refahi M, Sokhansanj BA, Mell JC, Brown JR, Yoo H, Hearne G, et al. Enhancing nucleotide sequence representations in genomic analysis with contrastive optimization. Communications Biology. 2025;8(1):517. 10.1038/s42003-025-07902-6.

[18] Refahi M, Hearne G, Muller H, Lynch K, Sokhansanj BA, Brown JR, et al. Fast and Scalable Gene Embedding Search: A Comparative Study of FAISS and ScaNN. In: Companion Proceedings of the 16th ACM International Conference on Bioinformatics, Computational Biology and Health Informatics. BCB Companion ’25. New York, NY, USA: Association for Computing Machinery; 2025. Available from: 10.1145/3768322.3769097.

[19] Boone J. Task geometry alignment enables parameter independent and accurate genomic search. Scientific Reports. 2026;16(1):24173. 10.1038/s41598-026-65239-4.

[20] Harrigan WL, Ferrell BD, Wommack KE, Polson SW, Schreiber ZD, Belcaid M. Improvements in viral gene annotation using large language models and soft alignments. BMC Bioinformatics. 2024;25(1):165. 10.1186/s12859-024-05779-6.

[21] Holur P, Enevoldsen KC, Rajesh S, Mboning L, Georgiou T, Bouchard LS, et al. Embed-Search-Align: DNA sequence alignment using Transformer models. Bioinformatics. 2025 03;41(3):btaf041. 10.1093/bioinformatics/btaf041. https://academic.oup.com/bioinformatics/articlepdf/41/3/btaf041/61778456/btaf041.pdf.

[22] Douze M, Guzhva A, Deng C, Johnson J, Szilvasy G, Mazaré PE, et al. The Faiss library. arXiv preprint arXiv:240108281. 2024;arXiv:2401.08281. [cs.LG].

[23] van Zyl DJ, Dunaiski M, Tegally H, Baxter C, de Oliveira T, Xavier JS, et al. Craft: a machine learning approach to dengue subtyping. Bioinformatics Advances. 2025 10;5(1):vbaf224. 10.1093/bioadv/vbaf224. https://academic.oup.com/bioinformaticsadvances/articlepdf/5/1/vbaf224/64518812/vbaf224.pdf.

[24] Marco-Sola S, Moure JC, Moreto M, Espinosa A. Fast gap-affine pairwise alignment using the wavefront algorithm. Bioinformatics. 2021 05;37(4):456–463. 10.1093/bioinformatics/btaa777. https://academic.oup.com/bioinformatics/articlepdf/37/4/456/50359789/btaa777.pdf.

[25] Marco-Sola S, Eizenga JM, Guarracino A, Paten B, Garrison E, Moreto M. Optimal gap-affine alignment in O(s) space. Bioinformatics. 2023 02;39(2):btad074. 10.1093/bioinformatics/btad074. https://academic.oup.com/bioinformatics/articlepdf/39/2/btad074/50530586/btad074.pdf.

[26] Li H. Minimap2: pairwise alignment for nucleotide sequences. Bioinformatics. 2018 05;34(18):3094–3100. 10.1093/bioinformatics/bty191.

[27] Holtgrewe M. Mason: A Read Simulator for Second Generation Sequencing Data. Institut für Mathematik und Informatik, Freie Universität Berlin; 2010. TR-B-10-06.

