## Additional file 1 for "Anniemap: Vector Search for Viral Short Read Alignment"

**Table 1** Recall of Bowtie2, BWA-MEM2 and Anniemap for different viruses across various read lengths at a standard error rate (2.5x mismatch scale).

| Virus | Bowtie2 | BWA-MEM2 | Anniemap |
|---|---|---|---|
| | | **50bp** | |
| CHIKV | 89.180 | 92.899 | 98.298 |
| DENV | 81.973 | 89.164 | 99.507 |
| Ebola | 97.276 | 98.764 | 99.845 |
| H1N1 | 87.812 | 88.776 | 96.247 |
| HCV | 34.234 | 36.453 | 57.978 |
| HIV | 55.665 | 62.018 | 87.251 |
| SARS-CoV-2 | 99.940 | 99.965 | 99.938 |
| | | **75bp** | |
| CHIKV | 95.612 | 97.070 | 98.587 |
| DENV | 92.794 | 97.366 | 99.931 |
| Ebola | 99.439 | 99.824 | 99.990 |
| H1N1 | 90.090 | 90.977 | 97.504 |
| HCV | 40.491 | 42.935 | 68.041 |
| HIV | 70.345 | 74.307 | 90.757 |
| SARS-CoV-2 | 99.979 | 99.982 | 99.995 |
| | | **100bp** | |
| CHIKV | 97.647 | 98.212 | 98.809 |
| DENV | 96.737 | 99.254 | 99.986 |
| Ebola | 99.855 | 99.980 | 100.000 |
| H1N1 | 91.369 | 92.099 | 98.375 |
| HCV | 44.479 | 46.331 | 77.875 |
| HIV | 77.826 | 79.203 | 95.431 |
| SARS-CoV-2 | 99.988 | 99.996 | 99.999 |
| | | **150bp** | |
| CHIKV | 98.607 | 98.712 | 99.027 |
| DENV | 98.993 | 99.926 | 99.995 |
| Ebola | 99.992 | 99.999 | 100.000 |
| H1N1 | 92.894 | 93.240 | 98.726 |
| HCV | 49.252 | 49.030 | 83.121 |
| HIV | 84.454 | 83.111 | 97.152 |
| SARS-CoV-2 | 100.000 | 100.000 | 100.000 |
| | | **300bp** | |
| CHIKV | 98.387 | 98.935 | 99.611 |
| DENV | 99.273 | 100.000 | 99.993 |
| Ebola | 99.997 | 100.000 | 100.000 |
| H1N1 | 94.050 | 95.034 | 98.682 |
| HCV | 51.768 | 51.181 | 84.614 |
| HIV | 88.754 | 86.830 | 98.069 |
| SARS-CoV-2 | 100.000 | 100.000 | 100.000 |

**Table 2** Recall of Bowtie2, BWA-MEM2 and Anniemap for different viruses across various read lengths at an extreme error rate (25x mismatch scale).

| Virus | Bowtie2 | BWA-MEM2 | Anniemap |
|---|---|---|---|
| | | **50bp** | |
| CHIKV | 44.558 | 54.485 | 93.259 |
| DENV | 36.570 | 45.256 | 90.873 |
| Ebola | 59.717 | 70.548 | 96.349 |
| H1N1 | 61.365 | 69.784 | 91.307 |
| HCV | 19.560 | 22.061 | 40.662 |
| HIV | 21.305 | 25.741 | 70.861 |
| SARS-CoV-2 | 79.166 | 88.013 | 94.067 |
| | | **75bp** | |
| CHIKV | 66.432 | 76.163 | 96.769 |
| DENV | 56.254 | 66.495 | 95.877 |
| Ebola | 80.507 | 88.790 | 99.091 |
| H1N1 | 76.843 | 82.512 | 93.592 |
| HCV | 26.207 | 28.576 | 48.839 |
| HIV | 35.312 | 41.199 | 77.733 |
| SARS-CoV-2 | 93.300 | 97.560 | 99.707 |
| | | **100bp** | |
| CHIKV | 77.213 | 85.337 | 98.296 |
| DENV | 67.327 | 77.637 | 98.291 |
| Ebola | 88.795 | 95.044 | 99.776 |
| H1N1 | 82.611 | 86.189 | 95.676 |
| HCV | 30.022 | 31.916 | 61.303 |
| HIV | 44.754 | 49.906 | 87.201 |
| SARS-CoV-2 | 97.394 | 99.376 | 99.960 |
| | | **150bp** | |
| CHIKV | 87.675 | 93.006 | 98.566 |
| DENV | 79.157 | 88.034 | 98.769 |
| Ebola | 95.661 | 98.831 | 99.887 |
| H1N1 | 87.085 | 88.370 | 96.331 |
| HCV | 34.482 | 35.031 | 67.535 |
| HIV | 56.604 | 58.442 | 89.742 |
| SARS-CoV-2 | 99.633 | 99.958 | 99.985 |
| | | **300bp** | |
| CHIKV | 90.868 | 97.792 | 99.133 |
| DENV | 85.043 | 97.992 | 99.717 |
| Ebola | 97.001 | 99.946 | 99.930 |
| H1N1 | 88.242 | 89.171 | 96.891 |
| HCV | 36.449 | 36.739 | 73.383 |
| HIV | 64.059 | 65.901 | 93.915 |
| SARS-CoV-2 | 99.667 | 99.999 | 99.994 |
